# Testing the basic tenet of the molecular clock and neutral theory by using ancient proteomes

**DOI:** 10.1101/821736

**Authors:** Tiantian Liu, Shi Huang

## Abstract

Early research on orthologous protein sequence comparisons by Margoliash in 1963 discovered the astonishing phenomenon of genetic equidistance, which has inspired the *ad hoc* interpretation known as the molecular clock. Kimura then developed the neutral theory and claimed the molecular clock as its best evidence. However, subsequent studies over the years have largely invalidated the universal molecular clock. Yet, a watered down version of the molecular clock and the neutral theory still reigns as the default model for phylogenetic inferences. The seemingly obvious tenet of the molecular clock on evolutionary time scales remains to be established by using ancient sequences: the longer the time of evolutionary divergence, the larger the genetic distance. We here analyzed the recently published Early Pleistocene enamel proteome from Dmanisi and found that ancient proteins were not closer to an outgroup than their orthologs from the extant sister species were. Together with a previous study, the combined results showed that most ancient proteins were in fact more distant to the outgroup. The results are unexpected from the molecular clock but fully predicted by the notion that genetic distances or diversities are largely at optimum saturation levels as described by the maximum genetic diversity (MGD) theory.

## Introduction

Since the early 1960s, protein and later DNA sequence comparisons have become widely used in building evolutionary trees [1-4]. Margoliash in 1963 discovered an astonishing finding, genetic equidistance where sister species are about equidistant to a simpler outgroup, and made an *ad hoc* interpretation of it by assuming a molecular clock [2, 5]. The molecular clock hypothesis assumes a constant and similar evolutionary rate among different species [1, 2, 5]. Thus, gene non-identity between species is thought to be largely a function of time. The molecular clock has been widely used in phylogenetic inferences and produced many controversial conclusions contradictory to phylogenies built by other independent methods, including the human relationship with the great apes and the origin of modern humans in Africa rather than Asia.[6-8].

The constant and similar mutation rate (i.e., molecular clock) interpretation of the equidistance result has not been verified by any independent observation and has on the contrary been contradicted by a large number of facts [9-15]. Nonetheless, researchers have treated the molecular clock as a genuine reality and have in turn proposed a number of theories to explain it [16-21]. The ‘Neutral Theory’ has found wide acceptance [19-21], even though it is widely acknowledged to be an incomplete explanation for the clock [13, 22], and an incomplete explanatory theory of nature in general [23-26]. The observed rate is measured in years but the Neutral theory predicts a constant rate per generation. Also, the theory predicts that the clock will be a Poisson process, with equal mean and variance of mutation rate. Experimental data have shown that the variance is typically larger than the mean.

Ohta’s “nearly neutral theory” explained to some extent the generation time issue by observing that large populations have faster generation times and faster mutation rates but remains unable to account for the great variance issue [27]. With the neutral and nearly neutral theory, molecular evolution has been treated as the same as population genetics or microevolution. However, the field still lacks a complete theory as Ohta and Gillespie had acknowledged [28]. The field has unfortunately yet to pay attention to the equidistance result, which has been considered by some as “one of the most astonishing findings of modern science”[29, 30].

While it is widely acknowledged that there is no universal molecular clock (vastly different species diverged for very long time do not have similar mutation rates), the seemingly obvious tenet of the molecular clock notion and the neutral theory, i.e., the longer the evolutionary divergence, the larger the genetic distance or sequence divergence, remains widely popular in phylogenetic inferences and has yet to be formally tested or established for species divergence over evolutionary time scales. A most direct, simple, and strait forward test of the molecular clock would be to use ancient proteins or DNAs from fossil species. Ancient fossil species are expected to show less sequence divergence from an outgroup than its extant sister species (Figure 1). Recent advances in protein sequencing methodology have made such test feasible. We here analyzed the recently published Early Pleistocene enamel proteome from Dmanisi [31] and found that ancient proteins are not closer to an outgroup than their orthologs from their extant sister species are. The results are unexpected from the molecular clock but are fully expected from a recently developed alternative framework.

**Figure 1.**
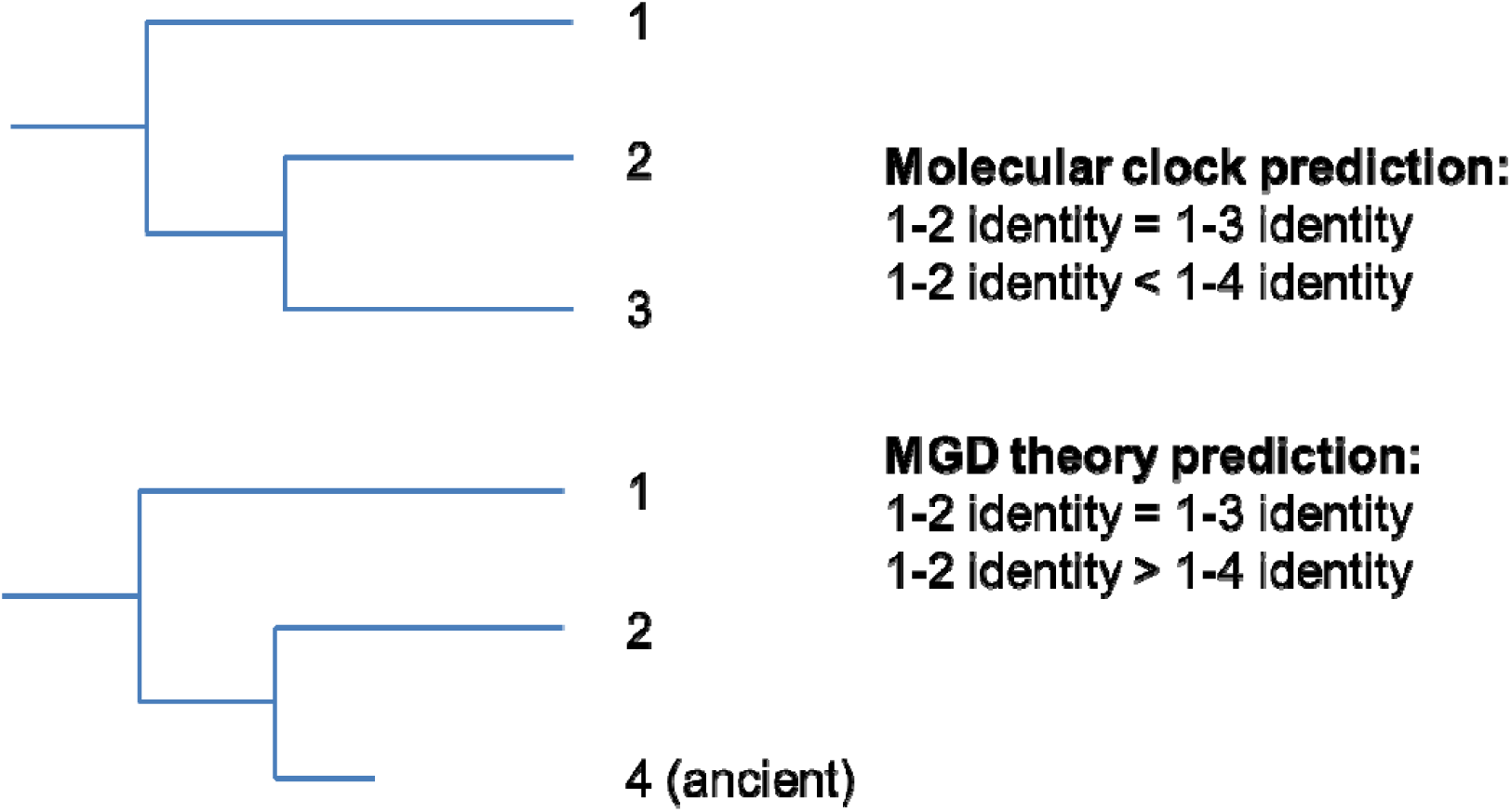
Schemes for testing the molecular clock hypothesis by using ancient proteins. Extant species are represented by numbers 1, 2, and 3. Ancient fossil species is represented by the number 4 with 2 representing the closest extant sister species of 4 and 1 representing the outgroup.

## Materials and Methods

Proteome sequences from Dmanisi were from the supplementary materials of the article by Cappellini [31]. Identification of closest extent proteins and comparisons with outgroups were performed using BLASTP against the protein database in Genbank. Extent proteins with the highest identity to the ancient samples were considered the closest extent proteins. Both the ancient and the extent orthologs were then aligned with the outgroup protein to determine which was closer to the outgroup.

## Results

The recently published Early Pleistocene enamel proteome from Dmanisi had a total of 10 proteins from 6 different fossil samples [31]. The ancient species represented include one Equidae, one Rhinocerotidae, and four Bovidae. Some species such as Bovidae were represented by more than one sample and some proteins were sequenced from more than one sample. However, different peptide fragments were sequenced from different samples. Thus, each of the total 22 protein sequences was unique and different from others.

Using the ancient sequences, we searched the Genbank to identify the orthologs from the closest extant sister species. We then compared both the ancient and the extant proteins to an outgroup. We first tested human as the outgroup to determine which shared more identity with human. The peptide fragments of each protein were each individually examined for identities between the outgroup and the sister species consisting of the ancient species and it closest extant species. Gaps were not counted and the same length and region of alignment were maintained for the ancient and its extant sister species. The results from all peptide fragments of a protein were then combined to obtain the total number of identical residues and the total alignment length (Table 1, and Supplementary Table 1). For a total of 22 proteins, five among them were too short to be informative in terms of revealing any differences between the two sisters. Of the remaining 17 proteins, fifteen showed lower number of identical residues between the ancient samples and the outgroup relative to between the extant proteins and the outgroup while two showed the opposite, indicating that the ancient proteins were significantly more distant to the outgroup human than the extant proteins were (P<0.01, Chi squared test).

**Table 1.** Identities between human as the outgroup and the ancient or present day species.

| Sample | Protein | Length | Identities to *Homo sapiens* |  |  |
| --- | --- | --- | --- | --- | --- |
|  |  |  | Ancient | Present | Difference |
| 16857 | COL1A2 | 613 | 546 | 554 | -8 |
| 16639 | AMELX | 89 | 70 | 77 | -7 |
| 16635 | ENAM | 249 | 184 | 189 | -5 |
| 16635 | AMELX | 173 | 146 | 150 | -4 |
| 16857 | COL3A1 | 507 | 385 | 389 | -4 |
| 16632 | AMELX | 139 | 117 | 120 | -3 |
| 16632 | MMP20 | 130 | 109 | 112 | -3 |
| 16635 | AMBN | 118 | 102 | 104 | -2 |
| 16635 | AMTN | 19 | 14 | 16 | -2 |
| 16638 | AMELX | 50 | 41 | 43 | -2 |
| 16635 | MMP20 | 57 | 48 | 49 | -1 |
| 16638 | AMBN | 139 | 119 | 120 | -1 |
| 16639 | AMBN | 115 | 103 | 104 | -1 |
| 16641 | AMBN | 92 | 82 | 83 | -1 |
| 16641 | AMELX | 59 | 54 | 55 | -1 |
| 16632 | ALB | 27 | 20 | 20 | 0 |
| 16632 | ENAM | 182 | 147 | 147 | 0 |
| 16638 | MMP20 | 44 | 42 | 42 | 0 |
| 16639 | ENAM | 123 | 98 | 98 | 0 |
| 16641 | ENAM | 67 | 57 | 57 | 0 |
| 16635 | ALB | 106 | 77 | 76 | 1 |
| 16638 | ENAM | 126 | 106 | 105 | 1 |

To verify the above results, we next tested a different outgroup *Sus scorfa*. Of the 24 proteins, four showed no difference between the sisters and hence were non informative due to probably the short length covered; 15 showed lower number of identical residues between the ancient samples and the outgroup relative to between the extant proteins and the outgroup and 3 showed the opposite (Table 2), indicating that the ancient proteins were significantly more distant to the outgroup *Sus scorfa* than the extant proteins were (P<0.01, Chi squared test).

**Table 2.**
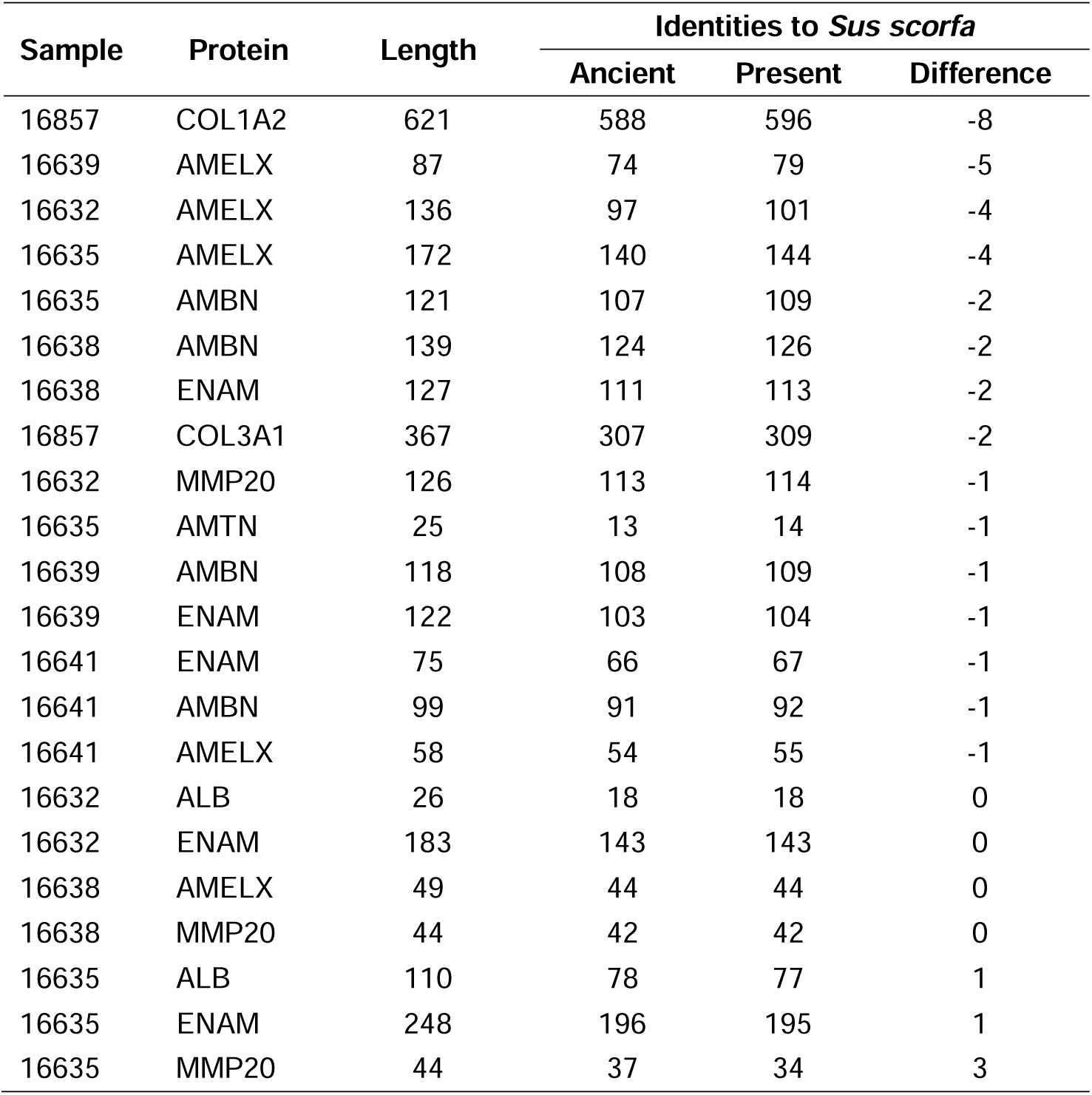
Identities between *Sus scrofa* as the outgroup and ancient or present day species.

| Sample | Protein | Length | Identities to *Sus scrofa* |  |  |
| --- | --- | --- | --- | --- | --- |
|  |  |  | Ancient | Present | Difference |
| 16857 | COL1A2 | 621 | 588 | 596 | -8 |
| 16639 | AMELX | 87 | 74 | 79 | -5 |
| 16632 | AMELX | 136 | 97 | 101 | -4 |
| 16635 | AMELX | 172 | 140 | 144 | -4 |
| 16635 | AMBN | 121 | 107 | 109 | -2 |
| 16638 | AMBN | 139 | 124 | 126 | -2 |
| 16638 | ENAM | 127 | 111 | 113 | -2 |
| 16857 | COL3A1 | 367 | 307 | 309 | -2 |
| 16632 | MMP20 | 126 | 113 | 114 | -1 |
| 16635 | AMTN | 25 | 13 | 14 | -1 |
| 16639 | AMBN | 118 | 108 | 109 | -1 |
| 16639 | ENAM | 122 | 103 | 104 | -1 |
| 16641 | ENAM | 75 | 66 | 67 | -1 |
| 16641 | AMBN | 99 | 91 | 92 | -1 |
| 16641 | AMELX | 58 | 54 | 55 | -1 |
| 16632 | ALB | 26 | 18 | 18 | 0 |
| 16632 | ENAM | 183 | 143 | 143 | 0 |
| 16638 | AMELX | 49 | 44 | 44 | 0 |
| 16638 | MMP20 | 44 | 42 | 42 | 0 |
| 16635 | ALB | 110 | 78 | 77 | 1 |
| 16635 | ENAM | 248 | 196 | 195 | 1 |
| 16635 | MMP20 | 44 | 37 | 34 | 3 |

The ancient sequences analyzed above had all isoleucines converted into leucines, as standard tandem mass spectrometry (MS/MS) cannot differentiate between these two isobaric amino acids. Leucines are about 2 fold more frequent than isoleucines in mammalian proteins. Thus, some leucines in the above analyzed ancient samples may in fact be isoleucines and would cause artificial non-identities with the outgroup. We next focused only on peptides that showed non-identities between the ancient samples and the sister taxon in amino acid positions not involving leucine/isoleucines. Three of five informative proteins showed lower identity between the ancient and the outgroup human relative to that between its extant sister taxon and human while two showed the opposite. With *Sus scorfa* as the outgroup, two of four informative proteins showed lower identity between the ancient and the outgroup relative to that between its extant sister taxon and the outgroup while two showed the opposite (Table 3). As the number of informative proteins were limited, the results were inconclusive with regard to ancient proteins being more distant to the outgroup but did show such a trend. At least, there was no indication that the ancient proteins were closer to the outgroup, as predicted by the molecular clock. Together with a previous study of ours that showed four informative ancient proteins to be all more distant to the outgroup [10], the combined studies showed that 7 or 6 ancient proteins were more distant to the outgroup while 2 were closer. Overall, these results were not expected by the molecular clock hypothesis.

**Table 3.**
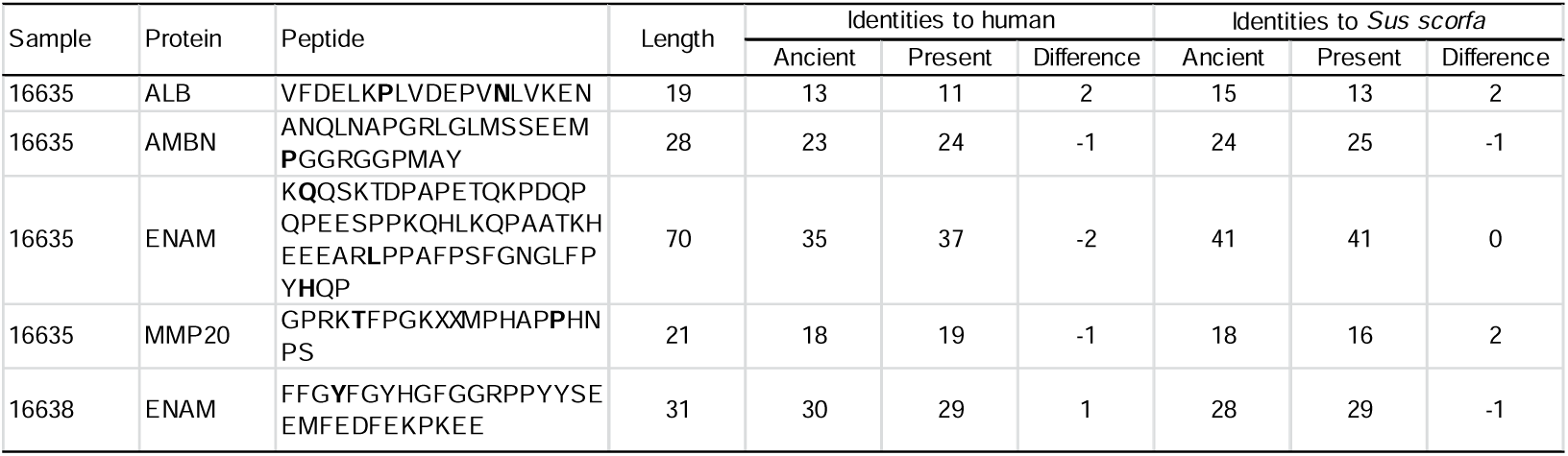
Identities between the outgroup and ancient or present day species. Amino acids differing between the ancient and its extant sister taxon are in bold.

## Discussion

Our results here showed that ancient proteins were not closer to an outgroup than their orthologs from the closest extant species of the ancient taxon, confirming a previous study of ours using ancient proteins [10]. These results remained even when uncertain amino acid positions due to technical limitations such as leucines/isoleucines were excluded from the analyses. The ancient proteins used here were from ∼2 million years ago and those from the previous study included a dinosaur protein from 68 million years ago. The timeframe concerned is thus long enough for ancient proteins to have lower number of substitutions than their orthologs from their extant sister taxons. And yet 7 or 6 ancient proteins were more distant to the outgroup while only 2 were closer. The results therefore were unexpected if the basic tenet of the molecular clock and neutral theory is true that non-identities in sequences is largely a function of time. That most of the ancient proteins were more distant to the outgroup was even more unexpected from the molecular clock and neutral theory. What then may explain the seemingly unexpected observations here?

In recent years, a more complete molecular evolutionary theory, the maximum genetic distance or diversity (MGD) hypothesis, has been making steady progress in solving both evolutionary and contemporary biomedical problems [6, 8, 25, 32-41]. That reality is largely at maximum genetic diversity or distance no longer changing with time is *a priori* expected and supported by numerous facts [12, 25, 42, 43]. Genetic distance can increase with time up to a point when maximum saturation is reached. At such maximum/optimum saturation, genetic distance would no longer be related to time but are determined by physiological selection and environmental selection. The MGD theory has solved the two major puzzles of genetic diversity, the genetic equidistance phenomenon and the much narrower range of genetic diversity relative to the large variation in population size [25, 26]. The primary determinant of genetic diversity (or more precisely MGD) is species physiology with complex physiology being compatible only with lower levels of random genetic noises/diversities [25, 44]. The genetic equidistance result of Margoliash in 1963 is in fact the first and best evidence for maximum distances rather than linear unsaturated distances as mis-interpreted by the molecular clock and in turn the neutral theory [2, 6, 11, 23, 25, 43]. Two contrasting patterns of the equidistance result have now been recognized, the maximum and the linear [6, 23]. The neutral theory explains only the linear pattern, which however represents only a minority of any genome today. The link between traits/diseases and the amount of SNPs shows an optimum/maximum (Pareto optimum) genetic diversity level maintained by selection, thereby providing direct experimental disproof for the neutral assumption for common SNPs [32-38]. More direct functional data invalidating the neutral assumption have also been found [45].

The results here are fully predicted by the MGD theory (Figure 1). If the observed genetic distance or non-identities of today and of the relatively recent past such as 2 million years ago as in the case of the Dmanisi samples here were at maximum saturation, it would not be determined by time but by physiology and environmental adaptations. As the ancient taxon and its closest present day sister species would be very close in physiology, their genomes would be highly similar if not identical in those parts involved in physiology. However, their genomes would be different in the parts involved in adaptation to environments since environmental conditions from ∼2 million years ago are expected to be different from today. As the outgroup species are from the present day time, they are expected to share some adaptive variants with all present day species as a common adaptive strategy to today’s environment. Hence, the sister species of the ancient taxon would be expected to be closer to the outgroup than the ancient is. The degree of this closeness would be related to the similarities between the ancient environments and today’s.

Using ancient protein or DNA sequences can be very effective in testing evolutionary theories. Our results here show that the basic tenet of the molecular clock and neutral theory does not hold for evolutionary time scales when maximum mutation saturation has been reached. This conclusion is expected to be further confirmed when more ancient sequences become available in the future.

## Acknowledgments

Supported by the National Natural Science Foundation of China grant 81171880 and the National Basic Research Program of China grant 2011CB51001.

## References

[1] Zuckerkandl E, Pauling L. Molecular disease, evolution, and genetic heterogeneity, Horizons in Biochemistry. New York: Academic Press, 1962.

[2] Margoliash E. Primary structure and evolution of cytochrome c. Proc. Natl. Acad. Sci. 1963;50:672–9.

[3] Doolittle RF, Blombaeck B. Amino-Acid Sequence Investigations Of Fibrinopeptides From Various Mammals: Evolutionary Implications. Nature 1964;202:147–52.

[4] Fitch WM, Margoliash E. Construction of phylogenetic trees. Science 1967;155:279–84.

[5] Kumar S. Molecular clocks: four decades of evolution. Nat Rev Genet 2005;6:654–62.

[6] Huang S. Primate phylogeny: molecular evidence for a pongid clade excluding humans and a prosimian clade containing tarsiers. Sci China Life Sci 2012;55:709–25.

[7] Johnson MJ, Wallace DC, Ferris SD, Rattazzi MC, Cavalli-Sforza LL. Radiation of human mitochondria DNA types analyzed by restriction endonuclease cleavage patterns. J Mol Evol 1983;19:255–71.

[8] Yuan D, Lei X, Gui Y, Zhu Z, Wang D, Yu J, et al. Modern human origins: multiregional evolution of autosomes and East Asia origin of Y and mtDNA. bioRxiv 2017:doi: https://doi.org/10.1101/106864.

[9] Huang S. Molecular evidence for the hadrosaur B. canadensis as an outgroup to a clade containing the dinosaur T. rex and birds. Riv. Biol. 2009;102:20–2.

[10] Huang S. Ancient fossil specimens are genetically more distant to an outgroup than extant sister species are. Riv. Biol. 2008;101:93–108.

[11] Huang S. The genetic equidistance result of molecular evolution is independent of mutation rates. J. Comp. Sci. Syst. Biol. 2008;1:092–102.

[12] Huang S. Inverse relationship between genetic diversity and epigenetic complexity. Nature Precedings 2008:doi.org/10.1038/npre.2009.1751.2.

[13] Pulquerio MJ, Nichols RA. Dates from the molecular clock: how wrong can we be? Trends Ecol Evol 2007;22:180–4.

[14] Nei M, Kumar S. Molecular evolution and phylogenetics. New York: Oxford University Press, 2000.

[15] Ayala FJ. Molecular clock mirages. BioEssays 1999;21:71–5.

[16] Van Valen L. Molecular evolution as predicted by natural selection. J. Mol. Evol. 1974;3:89–101.

[17] Clarke B. Darwinian evolution of proteins. Science 1970;168:1009–11.

[18] Richmond RC. Non-Darwinian evolution: a critique. Nature 1970;225:1025–8.

[19] Kimura M. Evolutionary rate at the molecular level. Nature 1968;217:624–6.

[20] Kimura M, Ohta T. On the rate of molecular evolution. J. Mol. Evol. 1971;1:1–17.

[21] King JL, Jukes TH. Non-Darwinian evolution. Science 1969;164:788–98.

[22] Ayala FJ. Molecular clock mirages. BioEssays 1999;21:71–5.

[23] Hu T, Long M, Yuan D, Zhu Z, Huang Y, Huang S. The genetic equidistance result, misreading by the molecular clock and neutral theory and reinterpretation nearly half of a century later. Sci China Life Sci 2013;56:254–61.

[24] Kern AD, Hahn MW. The Neutral Theory in Light of Natural Selection. Mol Biol Evol 2018;35:1366–71.

[25] Huang S. New thoughts on an old riddle: What determines genetic diversity within and between species? Genomics 2016;108:3–10.

[26] Leffler EM, Bullaughey K, Matute DR, Meyer WK, Segurel L, Venkat A, et al. Revisiting an old riddle: what determines genetic diversity levels within species? PLoS Biol 2012;10:e1001388.

[27] Ohta T. Slightly deleterious mutant substitutions in evolution. Nature 1973;246:96–8.

[28] Ohta T, Gillespie JH. Development of Neutral and Nearly Neutral Theories. Theor Popul Biol 1996;49:128–42.

[29] Denton M. Evolution: a theory in crisis. Chevy Chase, MD: Adler & Adler, 1986.

[30] Denton M. Evolution, still a theory in crisis. Seattle, WA: Discovery Institute Press, 2016.

[31] Cappellini E, Welker F, Pandolfi L, Ramos-Madrigal J, Samodova D, Ruther PL, et al. Early Pleistocene enamel proteome from Dmanisi resolves Stephanorhinus phylogeny. Nature 2019;574:103–7.

[32] Zhu Z, Lu Q, Wang J, Huang S. Collective effects of common SNPs in foraging decisions in Caenorhabditis elegans and an integrative method of identification of candidate genes. Sci. Rep. 2015:doi:10.1038/srep16904.

[33] Zhu Z, Yuan D, Luo D, Lu X, Huang S. Enrichment of Minor Alleles of Common SNPs and Improved Risk Prediction for Parkinson’s Disease. PLoS ONE 2015;10:e0133421.

[34] Lei X, Yuan J, Zhu Z, Huang S. Collective effects of common SNPs and risk prediction in lung cancer. Heredity 2018:doi:10.1038/s41437-018-0063-4.

[35] He P, Lei X, Yuan D, Zhu Z, Huang S. Accumulation of minor alleles and risk prediction in schizophrenia. Sci Rep 2017;7:11661.

[36] Lei X, Huang S. Enrichment of minor allele of SNPs and genetic prediction of type 2 diabetes risk in British population. PLoS ONE 2017;12:e0187644.

[37] Gui Y, Lei X, Huang S. Collective effects of common SNPs and genetic risk prediction in type 1 diabetes. Clin Genet 2017;93:1069–74.

[38] Yuan D, Zhu Z, Tan X, Liang J, Zeng C, Zhang J, et al. Scoring the collective effects of SNPs: association of minor alleles with complex traits in model organisms. Sci China Life Sci 2014;57:876–88.

[39] Yuan D, Huang S. Genetic equidistance at nucleotide level. Genomics 2017:10.1016/j.ygeno.2017.03.002.

[40] Luo D, Huang S. The genetic equidistance phenomenon at the proteomic level. Genomics 2016;108:25–30.

[41] Biswas K, Chakraborty S, Podder S, Ghosh TC. Insights into the dN/dS ratio heterogeneity between brain specific genes and widely expressed genes in species of different complexity. Genomics 2016;108:11–7.

[42] Huang S. Histone methylation and the initiation of cancer, Cancer Epigenetics. New York: CRC Press, 2008.

[43] Huang S. The overlap feature of the genetic equidistance result, a fundamental biological phenomenon overlooked for nearly half of a century. Biological Theory 2010;5:40–52.

[44] Romiguier J, Gayral P, Ballenghien M, Bernard A, Cahais V, Chenuil A, et al. Comparative population genomics in animals uncovers the determinants of genetic diversity. Nature 2014;515:261–3.

[45] Pontis J, Planet E, Offner S, Turelli P, Duc J, Coudray A, et al. Hominoid-Specific Transposable Elements and KZFPs Facilitate Human Embryonic Genome Activation and Control Transcription in Naive Human ESCs. Cell Stem Cell 2019;24:724–35 e5.

